# Perceptual constancy for an odour is acquired through changes in primary sensory neurons

**DOI:** 10.1101/2023.11.17.567529

**Authors:** Mark Conway, Merve Oncul, Kate Allen, Jamie Johnston

## Abstract

The ability to consistently recognise an object despite sensory input that varies with environmental conditions and/or distance from the object is termed perceptual constancy. This is not an innate ability, rather it develops early in life and is likely dependent upon experience (*1*, *2*). The neural mechanisms underpinning the development of perceptual constancy are poorly understood. We have taken advantage of the olfactory system of mice and show that when mice are naïve to an odour a perceptual shift occurs with increasing concentration. The perceptual shift coincides with a rapid reduction in activity of a single olfactory receptor channel that is most sensitive to the odour. This drop in activity is not a property of circuit interactions within the olfactory bulb, rather it is due to a sensitivity miss-match of olfactory receptor neurons within the nose. We show that after forming an association of this odour with food, the sensitivity of the receptor channel is matched to the odour object, preventing transmission failure and promoting perceptual stability. These data show that plasticity of the primary sensory organ enables learning of perceptual constancy.

---

The properties of objects are largely invariant yet they evoke different neural activity within primary sensory organs depending on, for example, their proximity to the sensor. Perceptual constancy, when the perception of an object remains the same despite changing neural input, arises early in life and is dependent upon experience (*1*, *2*). However, there are examples of perceptual changes resulting from intensity differences (*3–5*). The neural mechanisms that both give rise to intensity induced changes in perception and the mechanisms that promote perceptual constancy are poorly understood. We have taken advantage of the olfactory system of mice, where, in a laboratory setting, experience of odours are intrinsically restricted allowing us to compare experience induced changes in perception.

## Results

### Concentration dependent shifts in odour perception

To evaluate whether mice experience a perceptual change in response to varying concentrations of an odourant, we employed a cross-habituation assay, a standard method used to determine a rodent’s ability to differentiate between odourants (*6–10*). We used an automated approach based on (*11*), where mice were placed in a test chamber with odours delivered through a nose-poke containing a beam break that logged investigation time (Figure 1A). As cross-habituation assays rely on both the mouse detecting the odour and choosing to investigate, we began by using 2-heptanone, a component of mouse urine (*12*), with the rationale that mice should investigate this odour if detected. Indeed mice investigated 2-heptanone at the lowest concentration tested (6 × 10^−7^ %) and then rapidly habituated to two subsequent presentations, this habituated state was maintained even with a 100 fold jump in concentration to 6×10^−5^ % (Figure 1B). When the concentration was increased 10,000 times above that of the original, the mice once more investigated the odour with a similar pattern of habituation to further stimuli (Figure 1B). This indicates that mice perceived a qualitative change in the odour between 6×10^−5^ % and 6×10^−3^ %, but did not between the two lowest nor the two highest cocentrations. We next used ethyl tiglate an odour to which the mice were naïve, in this case the mice failed to investigate for all concentrations up untill 6×10^−1^ % (Figure 1B). In this case, the data suggests that the mice perceived a qualitive difference between the concentrations of 6×10^−3^ % and 6×10^−1^ %. Esters are reported to have a neutral valence in mice (*13*), potentially explaining the lack of investigation between 6×10^−7^ - 6×10^−3^ %. However, the failure to investigate could merely reflect the inability of the mice to detect the lower concentrations. We therefore developed a method to measure the sensitivity of mice to novel odours that is independent of internal motivation. We head-fixed the mice on a treadmill (*14*) and, with video recording, tracked key facial features with deeplabcut (*15*) (Figure 1C). In both humans and rodents, detection of a novel stimulus results in pupil dilation (*16–20*) and we find that the lowest concentration of ethyl tiglate at 1×10^−7^ % there was a significant increase in the frequency content linked to sniffing/active exploration (Figure 1Eii & iii, n=6). These data indicate that mice can detect the lowest concentration of ethyl tiglate (6×10^−7^ %), it evokes pupil dilation and an increase in sniffing behaviour. Together these data indicate that mice can detect both 2-heptanone and ethyl tiglate at the lowest concentrations tested, and that with increasing concentration a perceptual shift occurs, between 6×10^−5^ and 6×10^−3^ % for 2-heptanone, and between 6×10^−3^ and 6×10^−1^ % for ethyl tiglate. We next sought to determine the neural basis for generating distinct percepts of the same molecule at different concentrations.

**Figure 1:**
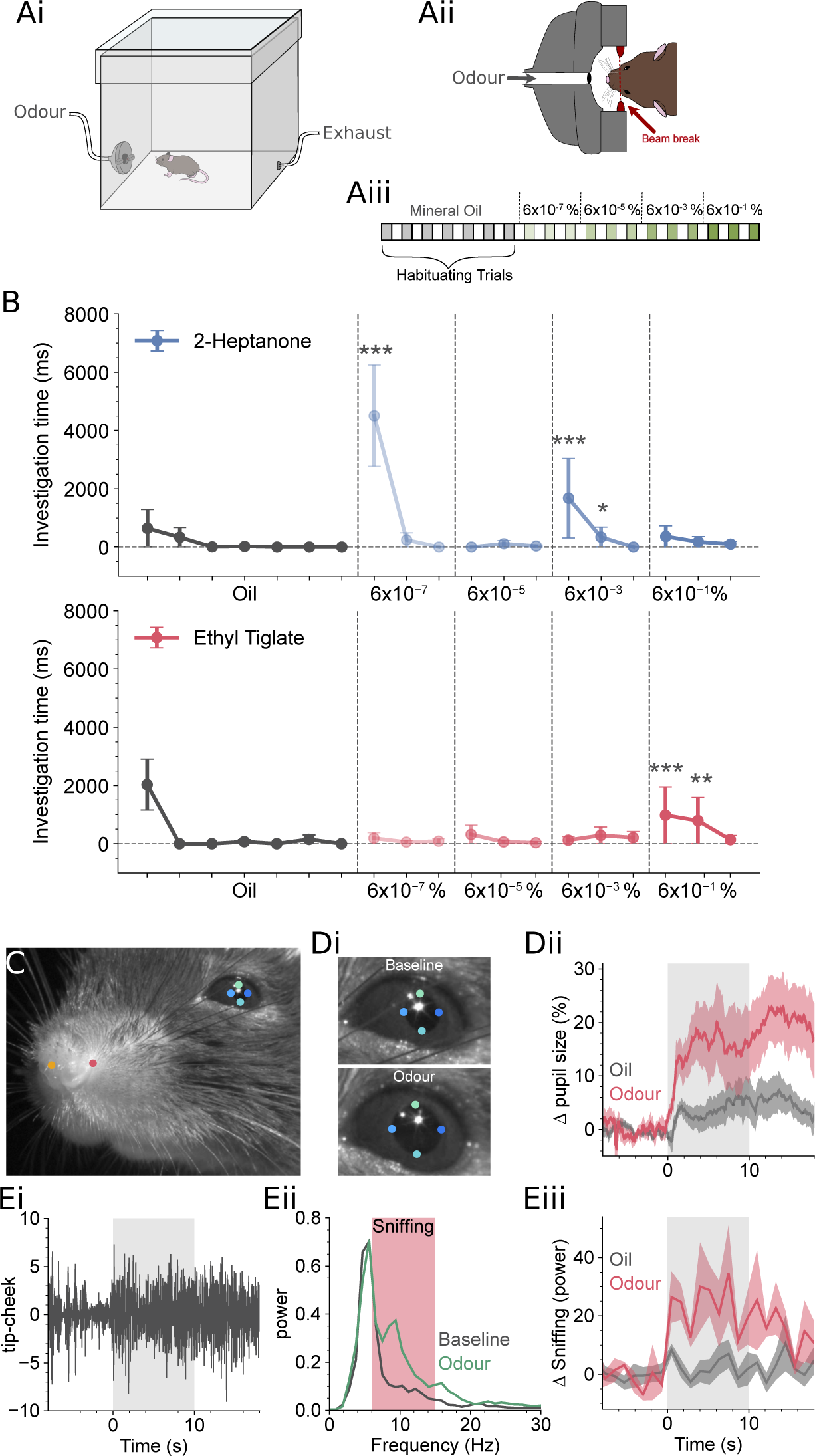
Measuring concentration-dependent changes in olfactory perception. Ai) Experimental paradigm, mice were placed in a test chamber with an odour delivery port and exhaust Aii) The odour delivery port contained a nose poke with beam break sensor to log investigation times. Aiii) Odour delivery protocol, each block represents 60 s (60 s stimulus, 60 s inter-stimulus interval). B) Odour investigation times during stimulus delivery for 2-heptanone and ethyl-tiglate, data are displayed as median ± the median absolute deviation, n=32. The horizontal dashed line indicates the basal amount of investigation calculated from the last 5 oil presentations. Asterisks indicate significance above the basal investigation rate. Odour concentrations are displayed as the final concentration measured at the nose poke. C) Mice were head-fixed and facial features were tracked with deeplab cut (see methods), coloured dots indicate key points tracked. Di) Pupil diameter before and during odour stimulation, diameter was calculated as the mean from the cardinal points. Dii) The relative change in pupil diameter displayed as mean ±SEM during presentaion of 1×10^−7^ % ethyl tiglate (red) and for 3 preceding stimulus blanks (grey) n=6, stimulus period indicated by shaded grey area. Ei) Oscillations in the distance between the key points for the nose tip and cheek. Eii) Fourier transforms of the data in Ei, for the 10 s before stimuli (grey) and during stimulation (green), sniffing band (7) indicated by shaded red box. Eiii) Change in the power for the sniffing band displayed as mean ±SEM during presentation of 1×10^−7^ % ethyl tiglate (red) and for 3 preceding stimulus blanks (grey) n=6, stimulus period indicated by shaded grey area.

### Odour percepts rely on a sparse code

To explore how the brain represents the range of concentrations used in Figure 1, we employed *in vivo* 2-photon imaging. We used the genetically encoded Ca^2+^ indicator GCaMP6f (*21*) expressed in mitral and tufted cells of the olfactory bulb, driven by the Pcdh21 promoter (*22*) (Pcdh21xGCaMP6f mice, see methods). Mitral and tufted cells form the output of the olfactory bulb and receive direct input from the olfactory nerve on their tuft dendrites located within a single glomerulus (*23–25*). We began by imaging the odour-evoked responses in the glomerular layer, the site of the initial excitation of these output neurons. This approach enables visualisation of the spatiotemporal activity arriving in the olfactory bulb (*26*), as each glomerulus corresponds to input from a single olfactory receptor (*27*). We presented mice with concentrations of ethyl tiglate spanning the entire range used in the cross-habituation experiments (Figure 1). We generated response maps (Figure 2A) by averaging glomerular activity over the 3 s stimulus period. As in the cross-habituation experiments, mice were presented with the most dilute concentration first, with each successive stimulus 3-10 fold stronger. Glomerular responses were detected at every concentration presented, supporting the finding that mice can detect ethyl tiglate over 6 orders of magnitude (Figure 1). In accordance with recent reports (*28*), glomerular responses to the weakest concentrations were sparse, with generally only a single glomerulus responding to the majority of concentrations presented from the weak percept (Figure 2A). As expected, the total number of active glomeruli was far greater when mice were presented with higher concentrations of the same odour, as has been reported previously (*29–31*). We assigned labels to the responses based on the cross-habituation experiment, responses between the weakest stimuli and ∼6×10^−3^ % were labelled as the ‘weak percept’ and responses above ∼6×10^−1^ % were labelled as the ‘strong percept’. We did not identify the precise concentration where the perceptual shift occurs, which may vary depending on nasal patency, but it falls between 6×10^−3^ and 6×10^−1^ % which we have termed the ‘transition range’ (Figure 2A).

**Figure 2:**
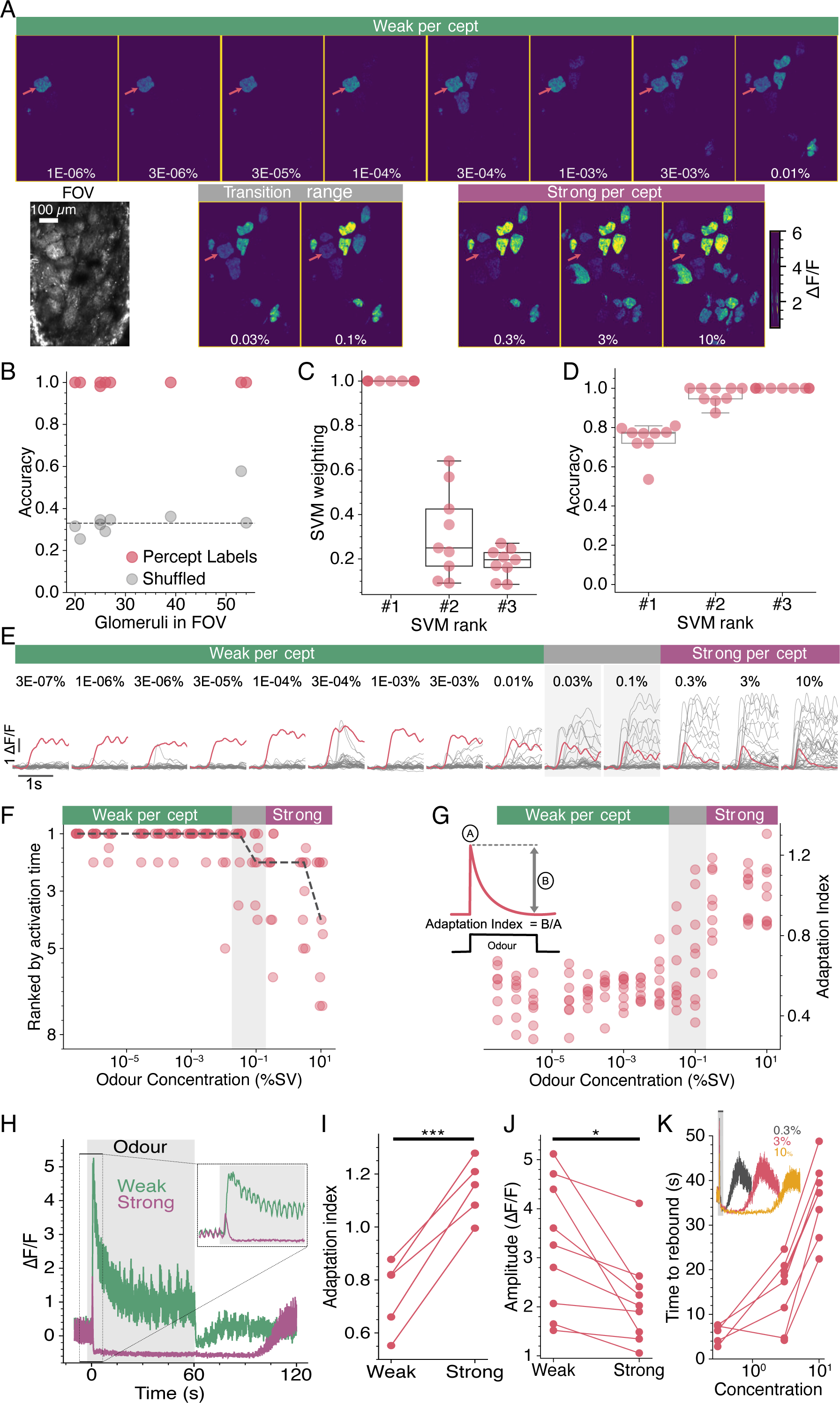
Neural correlates of perceptual shifts. A) Response maps and corresponding field of view in a Pcdh21xGCaMP6f mouse, showing the mean activity during 3 s odour stimuli for the concentrations indicated in white and grouped by odour percept (see text). Red arrows indicate the primary glomerulus. B) Linear SVM classifier performance using response maps from 9 mice (red dots). Classifier performance with shuffled labels (grey dots). C) The relative classifier weights for the top 3 glomeruli, the primary glomerulus has a weight of 1. D) Classifier performance using the top 3 glomeruli identified in B n=9. E) Responses of all glomeruli in A to single odour trials with the primary glomerulus shown in red, sampled at 42 Hz. Odour percepts indicated with coloured bars. F) The activation rank of the primary glomerulus as a function of concentration, each mouse represented by a dot, with jitter added for clarity. The dashed black line indicates the median n=9. G) The adaptation index of the primary glomerulus as a function of concentration, inset shows the calculation of adaptation index n=9. H) Response of a primary glomerulus to a 60 s stimulus of ethyl tiglate from the weak percept (1.0E^−6^ %) and strong percept (3 %). Inset shows expanded view of initial response and drop below baseline. Note the delayed rebound in activity long after the stimulus ends. I) The adaptation indexes of primary glomeruli to a 60 s stimulus grouped by percept n=5. J) Response amplitudes from the primary glomerulus for 3 s odour stimuli were larger for 3.0E^−3^ % (weak percept) than for 10 % (strong percept) n=9. K) The delay of the rebound in the primary glomerulus increases with stimulus strength, calculated from 3 s stimuli n=8, inset shows an example glomerulus with a grey bar indicating the 3 s stimulus.

Notably a linear classifier had a 99.8% success rate in predicting the odour percept based on the neural activity (Figure 2B, n=9). As the performance did not seem to depend on the number of glomeruli in the field of view we next examined the weights assigned to each glomerulus used in the classifier; these weights directly signify the extent to which each glomerulus contributes to the decision boundary. We found that a single glomerulus in each mouse made a major contribution, with the 2^nd^ and 3^rd^ most important having weights of 25 ±15% and 19 ±3% of the 1st (Figure 2C). Surprisingly, when we used only the single most important glomerulus the classifier achieved 74% accuracy and with only 2 glomeruli this increased to 96.7%, a comparable performance to using all glomeruli (Figure 2D). This suggests that only a few glomeruli are necessary to encode the odour percepts, rather than a broad pattern of active glomeruli.

Previous work has indicated that a sparse ‘primacy’ code may be used for odour identity (*32*, *33*), whereby the fastest activating glomeruli carry the most importance. Our data is consistent with such a primacy code; for each odour stimulus we determined the activation time of all responsive glomeruli (Figure 2E), when glomeruli were ranked in the order they activated we found that the glomerulus with the strongest predictive value was also the glomerulus that activated first (Figure 2E & F). However, this was only true for the weaker percept, for the strong percept this glomerulus began to lag behind other glomeruli that became active at higher concentrations (Figure 2E & F). Nevertheless, as this glomerulus contributed most to classifying the odour percept and was the first to activate for the weak percept we will refer to it as the ‘primary’ glomerulus. The most stricking behaviour of primary glomeruli was that they shift from a sustained response in the weak percept to rapid adaptation for the strong percept. We used the adaptation index (Figure 2G) to quantify the amount of adaptation as a function of concentration. An adaptation index (AI) of 1 indicates complete adaptation, whereas greater than 1 corresponds to adaptation that reduces the response to below baseline. As can be seen in Figure 2G, the amount of adaption of the primary glomerulus shifts from 0.34 ±0.03 for the highest concentration of the weak percept to near complete adaptation at the higher concentrations with an AI of 1 ±0.05 for the strong percept (p= 1.18×10^−7^, paired t-test n=9) and this shift to near complete adaptation occurs within the transition range (Figure 2G). Together these data indicate that odour percepts are likely generated using a sparse code, requiring just a few glomeruli and that a change in perception corresponds to rapid adaptation of the primary glomerulus.

The difference in how the primary glomerulus responds to weak and strong percepts becomes especially evident when 60 second stimuli are delivered (Figure 2H), mirroring the duration used in the cross-habituation experiments of Figure 1. Weak percepts show slow and incomplete adaptation (AI = 0.75 ±0.05 n=5), continuing to respond all throughout the stimulus, whereas, strong percepts generate rapid and complete adaptation. Strikingly, the response to stronger stimuli falls below baseline with an AI of 1.15 ±0.04 (Figure 2H & I, p = 0.0008, t-test, for the 5 animals where 60 s of both stimuli were delivered). Two further characteristics are of note when comparing responses to the strong and weak percepts; the peak amplitude was smaller for the stronger percept than the weak (Figure 2J, 2.03 ±0.54 vs 3.26 ±1.19 ΔF/F, p = 0.004, Wilcoxon, n = 9) and a rebound in activity was observed (1.35 ±0.28 ΔF/F, n=9), the delay to which depended on the strength of the stimulus (Figure 2K).

### Rapid adaptation is due to transmission failure from olfactory receptor neurons

What mechanism could give rise to rapid adaptation that generates a smaller peak response, an adapted response that falls below baseline and a subsequent rebound in activity? Such response dynamics are a hallmark of feed-forward inhibition (*34*, *35*), a circuit motif present at the glomerular layer (Figure 3A). The olfactory nerve input excites both the mitral/tufted dendrites—where the measurements in Figure 2 are taken—and inhibitory periglomerular neurons. Subsequently, the periglomerular neurons deliver delayed inhibitory drive to the mitral/tufted dendrites (*36*, *37*). To test whether such a mechanism gives rise to the fast adaption, we took advantage of mice where GCaMP6f expression is restricted to the olfactory receptor neurons (OMPxGCaMP6f, see methods) (*38*, *39*). If feedforward inhibition underpins the observed rapid adaption it should only be manifest in the mitral/tufted cells not in the olfactory nerve input. We were easily able to identify the same primary glomerulus in OMPxGCaMP6f mice as, across animals, glomeruli are located in almost identical locations (*40*) and at very low concentrations glomerular activation is sparse and structured (*28*) (Figure 2A & 3C). Surprisingly, the same phenomenon was evident in the olfactory nerve terminals of the primary glomerulus; when we compared 60 s responses between weak and strong percepts the same switch to rapid adaption was evident (Figure 3 D). The transition between sustained and adapting responses (Figure 3E) coincides with both the mitral/tufted transition (Figure 2G) and the perceptual switch (Figure 1B). This suggests that this phenomenon originates at the first synapse, and is not the result of post-synaptic processing. However, olfactory nerve terminals also receive GABA_B_ and dopamine D_2_ mediated feedback inhibition from periglomerular neurons, that act to reduce presynaptic calcium influx (*41–45*). To test whether feedback inhibition could explain the rapid adaptation in the olfactory nerve terminals we used topical application of CGP 54626 and raclopride, antagonists of GABA_B_ and D_2_ receptors respectively (*46*, *47*). Consistent with previous experiments (*48–50*), disrupting feedback inhibition led to increased pre-synaptic Ca^2+^ influx for both weak and strong percepts (Figure 3F-H), indicating that the drugs were exerting their expected action. However, with feedback inhibition disrupted the olfactory nerve terminals still displayed the same rapid adaptation to the strong percept (Figure 3F & I). These data demonstrate that rapid adaptation in the primary glomerulus, that coincides with a shift in odour percept, is not a feature that is computed by neural circuits in the brain, rather this signal is already present in the olfactory receptor neurons located in the nasal epithelium.

**Figure 3:**
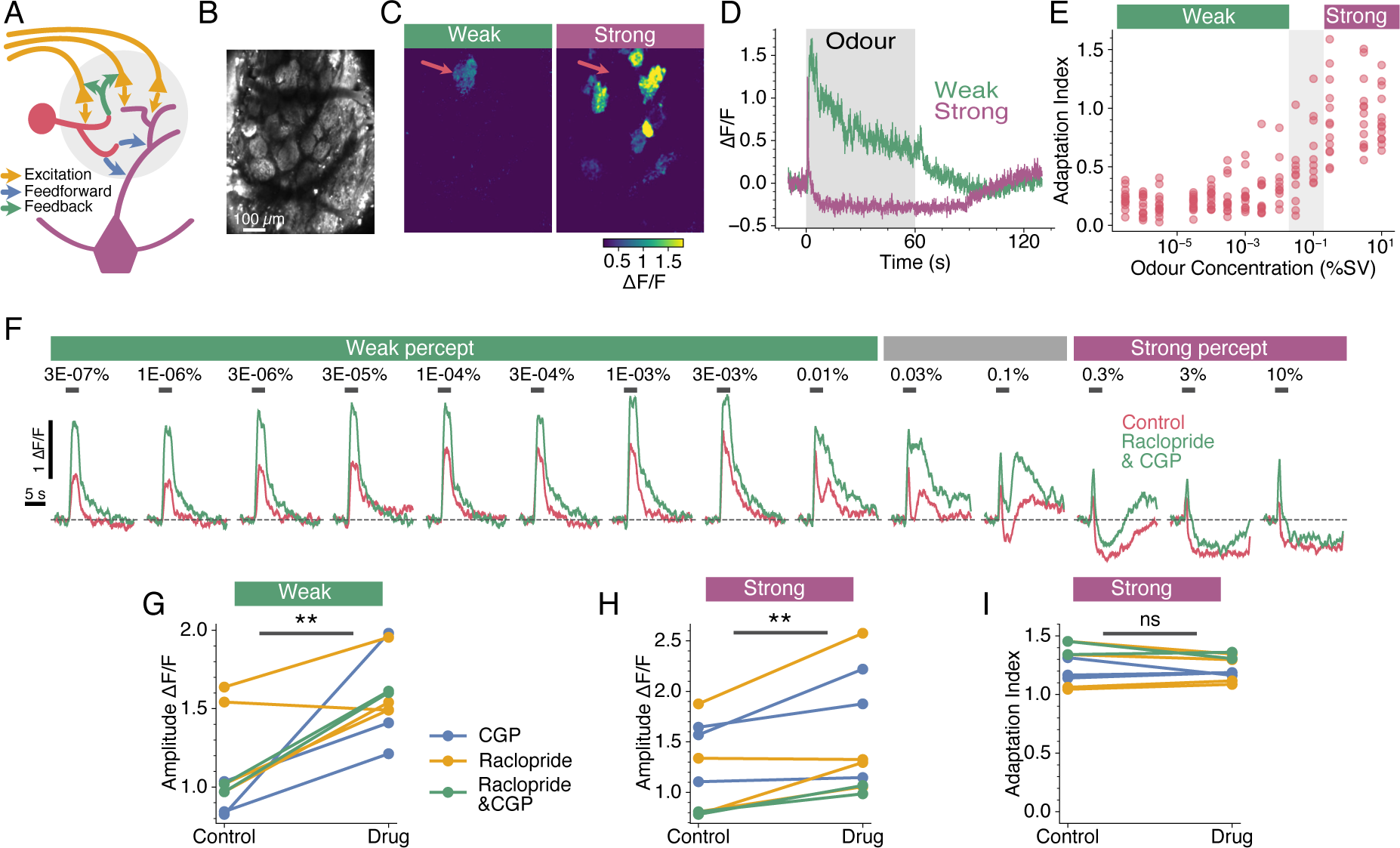
Rapid adaptation does not arise within the olfactory bulb. A) Intraglomerular circuitry within the olfactory bulb. Glutamatergic olfactory receptor neurons (yellow) synapse onto GABAergic periglomerular cells (red), prompting feedback and feedforward inhibition onto olfactory nerve terminals and mitral/tufted cells (purple), respectively. B) Field of view in an OMPxGCaMP6f mouse. C) Response maps for B, showing the mean activity during 60 s odour stimuli for a weak percept and strong percept (1.0E^−4^ % and 3 % respectively). Red arrows indicate the location of the primary glomerulus. D) Time courses of the responses in C. E) The adaptation index of the primary glomerulus increases when the concentration reaches the strong percept, calculated from 3 s stimuli n=13, calculation of adaptation index shown in Figure 2G). F) Responses of a primary glomerulus to single odour trials before and after application of the D_2_ and GABA_B_ antagonists raclopride and CGP 54626 respectively. G) & H) Response amplitudes from the primary glomerulus to 3 s odour stimuli before and after application of raclopride and CGP 54626 grouped by weak (G) and strong (H) percepts. Drug application enhances response amplitudes regardless of percept. Each ball and stick represents an individual mouse n=9. I) Adaptation indexes from the primary glomerulus to 60 s odour stimuli from the strong percept before and after application of raclopride and CGP. Note that drug application does not affect degree of adaptation n=9.

Adaptation of the olfactory transduction cascade has been well documented (*51–53*), where Ca^2+^ dependent feedback reduces sensitivity of the cyclic nucleotide gated current (*51*). However, it is hard to picture how such a mechanism could give rise to the adaptation that we observe in the olfactory nerve terminals, particularly as a decrease below baseline is observed during the stimulus (Figure 3D & I). To understand how this phenomena arises we employed a morphologically and biophysically realistic model of olfactory receptor neurons (Figure 4A). The model featured comparable membrane resistances and spontaneous spike rates as observed in *in vitro* recordings (*54*). To generate realistic receptor currents we constructed piecewise functions to reproduce reported receptor currents (*55*) (Figure 4B, see methods). In the model we could record the membrane potential from individual olfactory receptor neurons at both the soma and at the olfactory nerve terminals in response to receptor currents corresponding to weak and strong concentrations of odourant (Figure 4C & D). However, in our imaging experiments (Figures 2 & 3) we used a calcium indicator to measure the average activity due to the several thousand olfactory receptor neurons projecting to a glomerulus (*27*). To obtain equivalent recordings in our model we simulated 500 olfactory receptor neurons (Figure 4E) and convolved their mean spike rate with the kinetics of the GCaMP6f reporter (Figure 4F). This model provides important insight into the mechanism of rapid adaptation. The weak stimulus we provided had a peak receptor current of 13 pA and this resulted in a sustained increase in firing of individual olfactory receptor neurons, which showed slow adaptation at the population level (Compare Figure 4F with Figures 2E & 2H & 3D & 2F). In contrast, the strong stimulus we provided resulted in sustained depolarisation at the soma of the olfactory receptor neurons that generated a few action potentials at the onset of the stimulus that rapidly reduced in amplitude, due to accumulation of voltage-gated Na^+^ channel inactivation. The somatic membrane remained in a depolarised state preventing recovery from inactivation of the voltage-gated Na^+^ channels, thus blocking action potentials from passing down the axons. The resultant population spike rate, when convolved with the kinetics of the GCaMP6f reporter (Figure 4F), displays all the characteristics reported in Figures 2 & 3: 1) The brief initial burst of action potentials generates a smaller Ca^2+^ signal than the weaker stimulus, due to the low pass filtering of the GCaMP6f reporter (Figure 4F vs Figures 2H & J). 2) The response rapidly drops below the pre-stimulus baseline, due to the depolarising block terminating spontaneous action potential firing (Figure 2H, 3D and 4 E & F). 3) A rebound in action potential firing is observed after termination of the stimulus, as once the somatic membrane potential becomes sufficiently hyperpolarised to support recovery from inactivation the voltage-gated Na^+^ channels can resume generating action potentials (Figure 4C). We used a peak current of 96 pA for the strong stimulus, which is a rather conservative magnitude considering odour evoked receptor currents in rodents have been reported of >200 pA (*52*, *54*, *56*, *57*). Taken together, these data suggests that the shift in perception, occurring at higher concentrations, as depicted in Figure 1B, is a result of action potential failure within the primary sensory neurons situated in the nasal epithelium.

**Figure 4:**
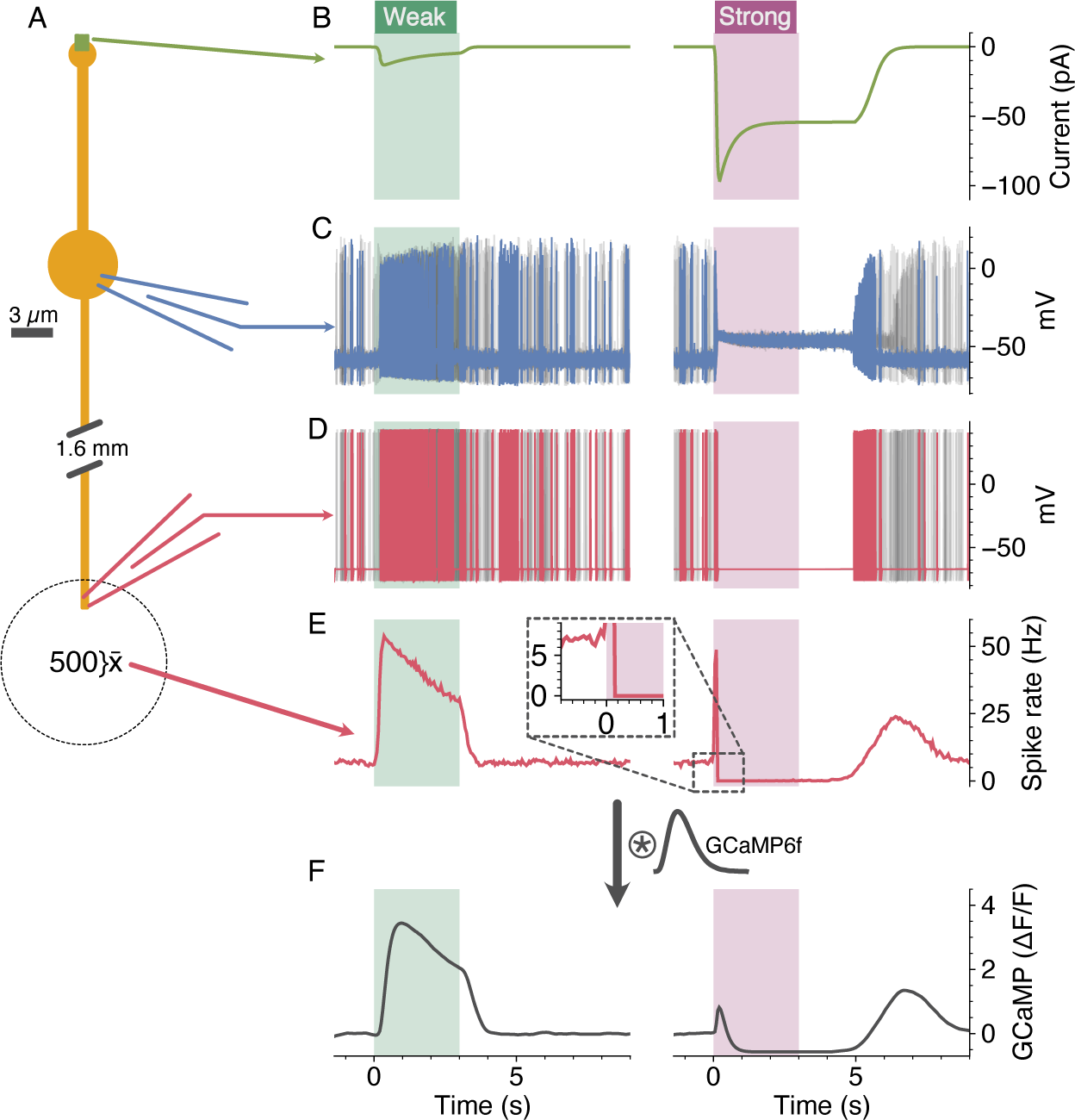
Depolarising block of olfactory receptor neurons results in rapid adaptation within the glomerulus. A) Morphology of model olfactory receptor neuron. B) Olfactory receptor currents for weak and strong odour stimuli. C) Somatic membrane potential recording for a single neuron in blue for weak and strong stimuli, with 4 further cells shown in grey. D) Axonal membrane potential recording for a single neuron in red for weak and strong stimuli, with 4 further cells shown in grey. E) Peri-stimulus time histograms showing the mean spike rates for 500 simulated neurons. Inset shows magnified view of the onset for the strong response, note the response falls below baseline. F) The spike rates from E convolved with the kinetics of the GCaMP6f reporter. See methods for model details.

### Learning perceptual constancy involves peripheral changes

We have described the failure of perceptual constancy for an unfamiliar odour and its underlying mechanism. The inability to recognise the same object across different concentrations would clearly be disadvantageous, particularly for salient odours such as food. Natural interaction with food stuffs would enable an animal to associate a range of concentrations with the same object, indeed, consuming the food provides much weaker activation of the olfactory epithelium by retronasal olfaction (*58*). We therefore investigated whether natural ‘passive’ association of the odour with food was able to endow perceptual constancy for ethyl tiglate across the full range of concentrations we used. We provided ethyl tiglate mixed with standard chow as a food source at a concentration corresponding to the strong percept (2.5 %). After 1 week of exclusively consuming ethyl tiglate scented food, we performed food finding tests. Mice, after an overnight fast, were placed in a cage with a buried food pellet, scented with 2.5% ethyl tiglate. Mice that had associated ethyl tiglate with food found the food pellet faster than a cohort of mice that had experienced the same amount of ethyl tiglate over the preceding 7 days but was not associated with food (154 ±96 vs 332 ±122 s, p = 0.036). The mice had clearly formed an association of ethyl tiglate with food as 11 of the 12 mice tested also began eating the pellet within the 10 min test, whereas only 1 of the 9 ‘exposed’ mice did so (Fig. 5A). These data indicate that the mice have formed an association of ethyl tiglate at the strong percept with food, we next sought to test whether this association extended to weaker concentrations that in naïve mice correspond to a different ‘weak percept’. Mice fed 2.5% ethyl tiglate were tested with a buried food pellet scented with 1×10^−3^ %. Remarkably all mice rapidly found the pellet and began eating (Fig. 5 A). We were concerned that, at this lower concentration, the smell of the standard chow may be aiding their localisation of the food, so we also performed the test with a buried cotton ball soaked in the same weak concentration, surprisingly all mice rapidly found and began nibbling the cotton ball (Fig. 5 A and supplemental video). These data indicate that after consuming a strong concentration of the odour mice associate a broad range of concentrations of the odour with food, even concentrations that previously evoked a different percept. We next asked what neural changes underpinned this learning induced change in perception; would the sensitive ‘primary glomerulus’ alter its properties to maintain responsiveness across the concentration range and/or shift to always being the first glomerulus to activate? Again we could easily identify the same primary glomerulus, it was the only one active at weak concentrations (Figure 5Bi). Remarkably, when we generated response maps similar to Figures 2A & 3C, the primary glomerulus was obvious at both the weak and strong concentrations (Figure 5B). This was due to the primary glomerulus responding throughout the whole stimulus period (Figure 5Bii), in stark contrast to what is observed in naïve mice (Figures 2H & 3D). This is quantified in Figure 5C which shows that the amount of adaptation in the primary glomerulus is much lower in mice that have associated ethyl tiglate with food (AI = 0.87 ±0.047, n=6) compared to naïve mice (AI = 1.14 ±0.028, p = 0.0021, t-test). We also examined the neural activity in mice that were exposed to the same amount of ethyl tiglate but not associated with food. In the majority of these ‘exposed’ mice the primary glomerulus still displayed close to complete adaptation with a mean AI of 1.002 ±0.045 across the 6 animals (Figure 5C & D), which was not significantly different to the naïve animals (p = 0.213, t-test). A shift from complete adaptation, caused by depolarising block, to a sustained response implies a shift in sensitivity of the primary glomerulus. Indeed, when we plotted the magnitude of response as a function of concentration a dramatic shift in sensitivity was observed between naïve animals and those that associated ethyl tiglate with food (Figure 5E), again with those only exposed to ethyl tiglate being intermediate between the two. Interestingly, the concentration at which the maximum response to ethyl tiglate occurred shifted by approximately two orders of magnitude, aligning closely with the concentration present in the food. This adjustment enables the primary glomerulus to effectively respond to the entire spectrum of concentrations that may be encountered during interactions with this particular food item. We expected that, after a food-association, the primary glomerulus would shift to being the first to activate across all concentrations, however it continued to lag behind other glomeruli at concentrations corresponding to the strong percept in naïve mice (Figure 5E). These data indicate that when mice make an association of an odour with food, changes must occur within olfactory receptor neurons that alter their sensitivity to the odour, this could be due to changes in the receptor compliment (*59*) or alterations of intrinsic membrane properties (*60*). However, the primacy (relative activation time) of the glomerular input appears to be unalterable, at least with natural interaction with an odour object.

**Figure 5:**
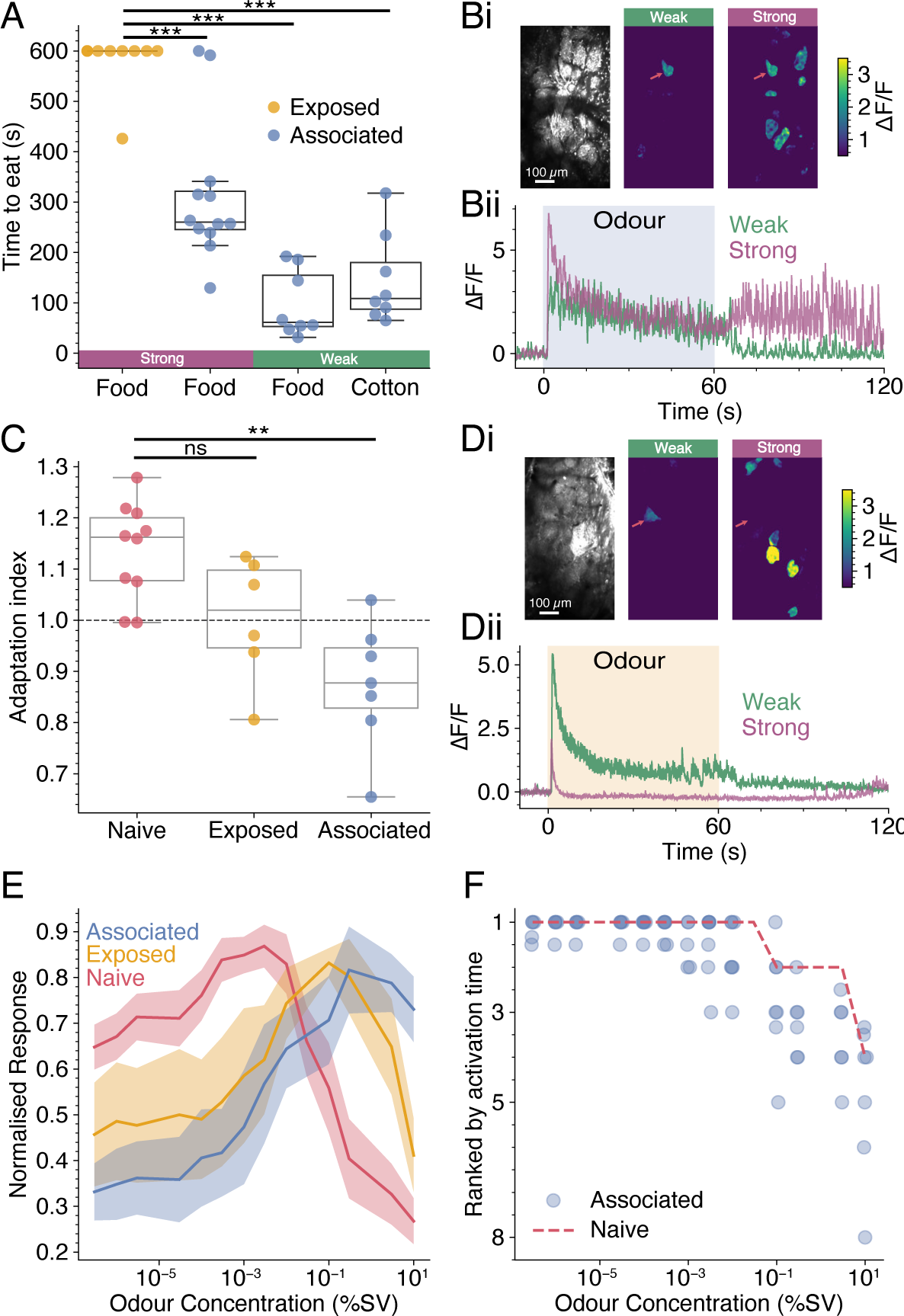
Learning perceptual constancy. A) Latency to eating for cohorts of mice either fed 2.5% ethyl tiglate mixed in their normal diet for 1 week (blue dots) or exposed to the same concentration (yellow dots). Mice were tested with a buried food pellet of the same concentration (purple bar) or with 1×10^−3^ % food pellet or cotton ball (green bar) Bi) Field of view and response maps for weak percept and strong percept (1×10^−3^ % and 3 % respectively) from a mouse after associating ethyl tiglate with food. Note the primary glomerulus is still evident in the strong percept. Bii) Time courses of the responses in Bi. C) The adaptation index for the primary glomerulus measured at the strong percept (3-10 %) for naïve, exposed and associated cohorts. D) Same as B but for mice that have been exposed to ethyl tiglate without food association. E) Normalised concentration response curves for the primary glomerulus displayed as mean ±SEM for the 3 cohorts. F) The activation rank of the primary glomerulus as a function of concentration, each mouse represented by a dot (n=7), the dashed red line indicates the median from naïve mice (n=9).

## Discussion

We show that mice experience a concentration-induced shift in odour perception (Figure 1), similar to reports in humans (*4*, *5*). This shift in perception coincides with a failure in transmission from receptor neurons to a single ‘primary’ glomerulus in the olfactory bulb (Figures 2-4). After association of the odour with food, such transmission failure is prevented and a single percept exists for a broad range of concentrations (Figure 5). These data are consistent with odour identity relying on a sparse code. Previous work has also suggested a sparse identity code based on the relative activation times of different glomeruli, with those activating earliest carrying more information (*32*, *33*). At weaker concentrations, and for naïve odours, our data are consistent with this model; the primary glomerulus activates first at weaker concentrations but at intensities perceived as a distinct percept it is no longer first (Figures 1B and 2F). However, after learning, when a broad range of concentrations evoke the same ‘food’ percept, the primary glomerulus at high concentrations still lags behind that of others despite obvious shifts in its sensitivity (Figure 5). It may be that the nature of the task may influence the coding strategy employed. When mice are trained in operant discrimination tasks, where reward is contingent upon a prompt behavioural action, mice learn to make discrimination decisions quickly (*32*, *33*), albeit after weeks of training. This is markedly different to what occurs with natural interaction with an odour object. In our experiments mice passively interact with odourised food, and by doing so will experience a range of concentrations of the odour that they will correlate with their distance to the object (*61*). In such a scenario the activity of the most sensitive glomerulus will show the highest degree of covariance with the odour stimuli, for example, it will be the only one activate at larger distances from the object. Over repeated interactions this primary glomerulus would therefore be given the most weight in determining the presence of the object. This idea of coherent covariation is used to explain the acquisition of semantic concepts (*62*) and would naturally give rise to a sparse odour identity code, especially with the observation that at low concentrations a sparse and structured representation of chemical space exists in glomerular activity (*28*). Such sparse codes for monomolecular odours are likely well suited to encoding the more complex mixtures found in natural odours, achieved by linearly combining the sparse representations of their individual constituents (*63*, *64*).

Our data show that the rapid adaptation underpinning a perceptual shift for the naïve odour is due to action potential failure within the olfactory receptor neuron (Figures 3 & 4), such behaviour at higher odour concentrations is evident in many recordings from olfactory receptor neurons (*53*, *56*, *65*). This failure of transmission occurs due to a miss-match between membrane resistance and receptor currents; olfactory receptor neurons have very high input resistances of ∼4-5 GΩ (*54*), whereas odour-evoked currents in these cells can be as large as 200 pA (*52*, *54*, *56*, *57*). With a simple minded ‘ohmic’ calculation, such receptor currents would cause a 800-1000 mV depolarisation. Of course the receptor current is not an ideal current source, rather it has a reversal potential, dominated by the Ca^2+^ activated Cl^−^ current ANO2 (*52*, *66*). So rather than a 800 mV depolarisation the membrane will become clamped at a depolarised potential. This sustained depolarisation locks voltage-gated Na^+^ channels in their inactivated state, preventing transmission of action potentials down the axon. This seems like a flaw in how the olfactory system operates, unless one considers that the primary goal of the olfactory system is first to detect odours and then to classify them. After exposure to a salient odour the olfactory receptor neurons adjust their sensitivity so that their maximum response falls near the concentration of the salient object and transmission failure no longer occurs at high intensities supporting perceptual stability (Figure 5). Such plasticity within the nose bears resemblance to aversive conditioning, whereby an odour paired with a foot shock brings about increased glomerular input for the conditioned odour by generating more olfactory receptor neurons carrying that receptor (*67*, *68*). It seems then that the nasal epithelium is a particularly dynamic structure able to tailor cell generation and receptor densities to optimally encode salient features encountered in the environment.

## Methods

### Animals

Animal handling and experimentation was carried out according to UK Home Office guidelines and the requirements of the United Kingdom (Scientific Procedures) Act 1986 and the University of Leeds animal welfare ethical review board. Mice were housed under a 12:12 h light/dark cycle with free access to food and water. All efforts were made to minimise animal suffering and the number of animals used. Pcdh21-nCre mice (C57BL/6Cr-Tg(Pcdh21-cre)BYoko (RBRC02189)), and OMP-Cre mice (B6;129P2(Cg)-Omp<tm4(cre)Mom>/MomTyagRbrc (RBRC02138)) were crossed with floxed GCaMP6f mice (GCaMP6f.flox, stock 028,865, B6J.CgGt(ROSA)26Sor < tm95.1 (CAGGaMP6f)), to generate Pcdh21xGCaMP6f mice, and OMPxGCaMP6f mice, respectively. Pcdh21-nCre and OMP-Cre mouse lines were originally obtained from RIKEN BioResource Research Center (Ibaraki, Japan), with permission from P. Mombaerts the original developer of the OMP-cre line (*38*, *39*). The GCaMP6f mouse line was obtained from Jackson Laboratory (Maine, USA). All mouse lines were maintained in house. Consistent with the NC3Rs guidelines (https://www.nc3rs.org.uk/who-we-are/3rs), both males and females aged 2-4 months old were used in this study.

### Odour stimuli

Odourants were obtained from Sigma-Aldrich, or Alfa Aesar. Liquid dilutions of odourants were prepared to achieve desired concentrations of approximately 3×10^−5^ %, 1×10^−4^ %, 3×10^−3^ %, 1×10^−2^ %, 0.1 %, 1 %, 3 % and 100 % using serial dilutions. Odourants were diluted in oil either (Sigma-Aldrich, 69794) or (Spectrum Chemical, C3465) within ∼1 week of experiments. Diluted odourants were delivered in vapour phase in synthetic medical air using either an 8 or 16 channel olfactometer (Aurora Scientific, 206A or 220A, respectively). Total flow rates from the olfactometers was kept constant at 1000 sccm. In imaging experiments, the output tubing of the olfactometer was positioned 1-2 cm in front of the mouse’s nose. Odourant presentations were always delivered in increasing concentrations. Inter-stimulus intervals were extended as odour concentration increased, varying between 20-120 s to minimise any adaptation. All odour concentrations are reported as % saturated vapour. Odour concentrations delivered to the behaviour boxes used in Figure 1 were measured with a miniPID (Aurora Scientific, 200B) placed at the nose port and are reported relative to the % saturated vapour used for imaging experiments.

### Cross-habituation test

Cross-habituation experiments were set up similar to the method described by (*11*). 2–3-months old mice were placed in a 25 x 25 cm perspex chamber with all sides opaque. Each chamber was fitted with an odour port and exhaust tube at opposing sides. The output of the olfactometer was connected to the odour ports of 4 chambers using identical path lengths of teflon tubing, the flow rate from the olfactometer was 1000 sccm. There was no difference in the concentration of odour delivered to each box as measured with a minPID. Each odour port housed an IR beam brake sensor (The Pi hut). Beam brake events and valve openings were logged using a MicroPython pyboard lite (v1.0) and pyControl GUI (v1.6). A mini vacuum pump (SLS2602) was attached to the exhaust tubes of all four chambers via tubing with an identical path length and air was extracted at a rate of 5.5 l/min. In each trial, mice were presented with either oil or a test odour for 60 s, followed by 60 s of synthetic medical air. Wild type C57bl6 mice were first habituated to the test environment for 10 minutes, before starting the stimulus protocol shown in Figure 1Aiii. Each presentation lasted 60 s with 60 s of medical air between presentations. In all instances, animals were naïve to the stimuli. Each animal was tested with 2-heptanone and ethyl tiglate in a pseudo-randomised order with 1 day between experiments.

### Head-fixed perception tests

Wild type C57bl6 mice were anaesthetised with isoflurane on a custom stereotaxic frame for head-bar attachment. Anaesthesia was maintained at a level of ∼1.5-2 % isoflurane, 1 l/min O2 during surgery. Metacam (5 mg / kg S.C.) and buprenorphine (0.1 mg / kg I.P.) were administered as analgesics. A small piece of skin above the skull, big enough for placing the head bar was carefully removed and cleaned with sterile saline solution. Superglue was initially applied over the exposed skull followed by dental cement to affix a custom 3D printed head bar. Additional dental cement was applied to cover the head bar and the exposed skull. Post surgery mice were given soaked diet and buprenorphine (0.1 mg / kg I.P.) for the following two days, all mice were allowed 1 week for recovery before habituation to head-fixation began. Mice were handled 5 minutes each day for 2 days prior to behavioural tests, aiming to acclimate them to the experimenter. Mice were head-fixed upon on a treadmill, described in (*14*), and habituated for 10 to 20 minutes per day for 2-3 days before recordings. The mouse face was imaged with a Basler camera (Cat. No: 107652) with 12 mm Edmund Optics lens (Cat. No: 33-303) and videos were captured at 120 Hz with 750 nm illumination (outside the visual range of mice). Odours were delivered using an olfactometer (220A, Aurora Scientific) and custom written code. The recording and synchronisation of data was performed with Bonsai-Rx (*69*) and a Teensy 4.2 microcontroller (PJRC). Each video acquisition was 35 s, composed of 10 s of baseline, 10 s stimulus and 15 s post stimulus. Each mouse was first presented with 5-7 oil trials before the the odour and all trials were spaced ≥ 60 s apart. A deeplabcut (*70*) neural net was trained on 15 frames from each mouse and used to extract the xy coordinates of the key points from every frame.

### Passive odour association

The diets of mice were supplemented with ethyl tiglate for 7 days. Ethyl tiglate was diluted in distilled water (1:40), before being combined with their regular diet in powdered form (equal W/V) and shaped into a single ball ( ∼ 5 g per mouse). Each mouse received a fresh food ball daily at ∼ 17:30 in a glass dish (7.5 cm W, 4.25 cm).

### Perceptual odour exposure

The environments of wild type mice were enriched with ethyl tiglate for 7 days. Ethyl tiglate was diluted in oil (1:40) and 1 ml was applied to Whatman Filter paper. Odourised filter paper was folded inside a metal tea ball and placed inside the animals home cage, replenished daily at ∼ 17:30.

### Food and odour finding test tests

Wild type mice were fasted for ∼ 16 hours before testing commenced. A clean housing cage was filled with ∼ 4 cm of fresh bedding, and an odourised food or cotton ball (∼ 1.5 cm^3^) was hidden beneath the bedding in a single corner. Care was taken not to leave odour trails during food/cotton ball placement. Mice were placed in the cage and a timer was set once a clear perspex lid had been attached. The time taken for mice to locate (defined as when the majority of the food/cotton ball became visible) and start eating the food/cotton ball was manually recorded.

### In vivo 2-Photon Ca^2+^ imaging

Mice were anaesthetised with urethane (1.5 g/kg) and body temperature was maintained at 37° C. Animals were secured with a custom made head bar and a craniotomy covering the right hemisphere of the olfactory bulb was performed. The exposed bulb was covered with 2 % low-melting point agarose in artificial cerebrospinal fluid and a 3 mm glass coverslip (Biochrom) was affixed with dental cement. Silicone rubber (Body double Fast Set) was applied to the skull surrounding the cranial window to create a well for the water dipping objective of the microscope. For experiments where drugs were topically applied, segments of dura were removed and the animal was imaged without a coverslip. GCaMP6f fluorescence was imaged with a custom built microscope, excited at 940 nm using a pulsed Mai Tai eHP DeepSee TI:sapphire laser system (SpectraPhysics). A resonant-galvo mirror assembly (Sutter instruments) scanned the beam through a 16 x water-dipping objective (N16XLWD-PF, NA 0.8, Nikon). Fluorescence was detected using GAasP photo-multiplier tubes and appropriate filters and dichroic mirrors. Images were acquired at 30-120Hz, using ScanImage software (*71*).

### Pharmacology

The GABA_B_-receptor antagonist CGP 54626 hydrochloride (Tocris Bioscience) was used at a concentration of 5 μM. The dopamine D_2_-receptor antagonist raclopride (Tocris Bioscience) was used at a concentration of 100 μM. Both drugs were dissolved in artificial cerebrospinal fluid (7.4 pH, 135 mM NaCl, 5.4 mM KCl, 5 mM HEPES, 1.8 mM CaCl_2_ 2H_2_O) and topically applied to the olfactory bulb 20 minutes before imaging recommenced.

## Data analysis

### Image segmentation of glomeruli

The Suite2p pipeline v0.10.1 (*72*) was used to register data with the default options (’nimg_init’: 300, ‘batch_size’: 500, ‘maxregshift’: 0.1, ‘smooth_sigma’: 1.15) regions of interest corresponding to glomeruli were manually drawn in FĲI (*73*) and raw fluorescence was extracted from glomeruli using custom-written code in Python. Extracted fluorescent traces were normalised as ΔF/F using the following equation: F-F^0^/F^0^, where F is the raw fluorescent trace and F^0^ is the baseline fluorescence recorded 5 s prior to odour stimuli.

### Adaptation index

To quantify the amount of adaptation we defined the adaptation index as the difference between the peak response (A in Figure 2G) and the mean of the last 100 ms of the stimulus period divided by the peak response. Prior to calculating AI data were filtered with a 5 point mean filter.

### Response maps

For each stimulus, response maps were generated using the following equation: F-F^0^/F^0^, where F is the raw fluorescent movie and F^0^ is the mean fluorescence recorded 3 s preceding the odour stimulus. Maps are displayed after 2D gaussian filtering with a sigma of 2 and areas outside the segmented glomeruli were set to zero.

### Classifier

We calculated the responses for each glomerulus by taking the mean of the Ca^2+^ signal over the stimulus period and 1 s after odour cessation (accounting for delayed activation seen in some glomeruli). Glomeruli were only considered to be responsive if their signal to noise ratio was ≥ 5, defined as: (mean amplitude over stimulus window - mean amplitude over 3 s preceding stimulus) / standard deviation over 3 s preceding stimulus, and that successive concentrations of the same odour were responsive. Trials where irregular breathing was apparent were excluded, i.e. a drop in activity across all glomeruli. 1-3 trials of each odour concentration were delivered and responses were normalised to the maximum response across all stimuli. Odour responses were assigned a percept label if they were within 50 % of the boundary concentration shown in Figure 1B. These data were classified using a linear support vector machine (class weight = balanced) from the scikit-learn library. The classifier accuracy was evaluated using the Leave-One-Out cross-validator to calculate weighted average F1 scores as reported in Figures 2B & D. Relative glomerular weighting (Figure 2C) were obtained by calculating the absolute values of the coefficients for each glomerulus and normalising each value to the largest assigned weight.

### Ranking glomerular activation times

To determine the first active glomerulus for a given stimulus, we identified the first frame during the stimulus period with a signal-to-noise ratio ≥ 5. The timestamp of this initial frame was taken as the activation time for the glomerulus. Trials where irregular breathing was apparent was excluded. For each stimulus, all responsive glomeruli were assigned a rank, with the first active glomerulus assigned a rank of 1. For trials with multiple repeats, the primary glomerulus was assigned the mean rank it received across all trials of the same concentration.

### Statistical analysis

For all statistical parameters data were first tested for normality with Shapiro Wilk and are reported as mean ± standard error of the mean if normal or median ± median absolute deviation if not. Paired comparisons employed a t-test or wilcoxon signed rank as appropriate. For the data in Figure 1B each of the investigation times for odour delivery was tested against pooled investigation times from the habituated oil (the last 5 oil presentations). All p values were adjusted for multiple comparisons with Bonferroni correction. When asterisks are used to indicate significance, * = p<0.05, ** = p<0.01 and *** = p<0.001.

## Model

Morphologically realistic models of olfactory receptor neurons and their receptor input were simulated in NEURON 8.2 (*74*, *75*). Each OSN consisted of 4 compartments: an axon of length 1.6 mm and diameter of 0.6 µm, a soma with diameter of 5 µm, a dendrite with a length of 12 µm and 0.8 µm diameter and an endbulb of 2 µm diameter. Axial resistance was 180 Ω * cm and membrane capacitance was 1 µF cm^−2^. Standard Hodgkin-Huxley channels were used at a uniform density throughout the cell with the following conductance densities: Na = 32 mS cm^−2^, K = 8 mS cm^−2^, passive = 0.02 mS cm^−2^, passive reversal −50 mV. This gave an input resistance of 4.6 GΩ similar to the reported membrane resistance of OSNs (*54*). To mimic the basal firing activity of olfactory receptor neurons evoked by spontaneous Nav channel openings in the cell body (*76*) gaussian noise with a mean of 1 pA and SD of 0.021 was injected into the somatic compartment which generated spontaneous firing at ∼4 Hz similar to the reported spontaneous rates (*54*). The receptor currents were modelled as a point process placed on the tip of the endbulb with a time course described by 3 piecewise functions obtained from fits to the synaptic currents reported in (*53*). The 3 piecewise functions correspond to the onset and duration of the odour stimulus (a), the decay after the stimulus (b) and the adaptation during the steady-state phase of the stimulus (c). The synaptic conductance (g) was therefore g = m(a+b-c) where m is a scaling factor. For the weak odour concentration:

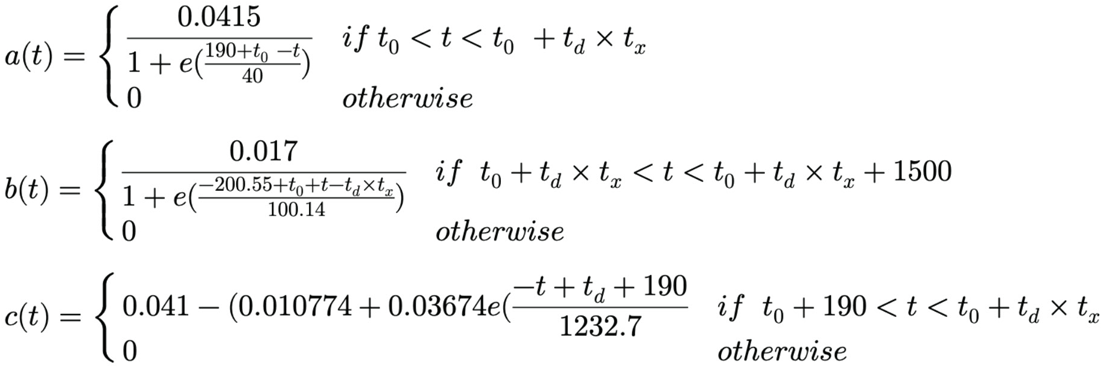

And for the strong odour concentration:

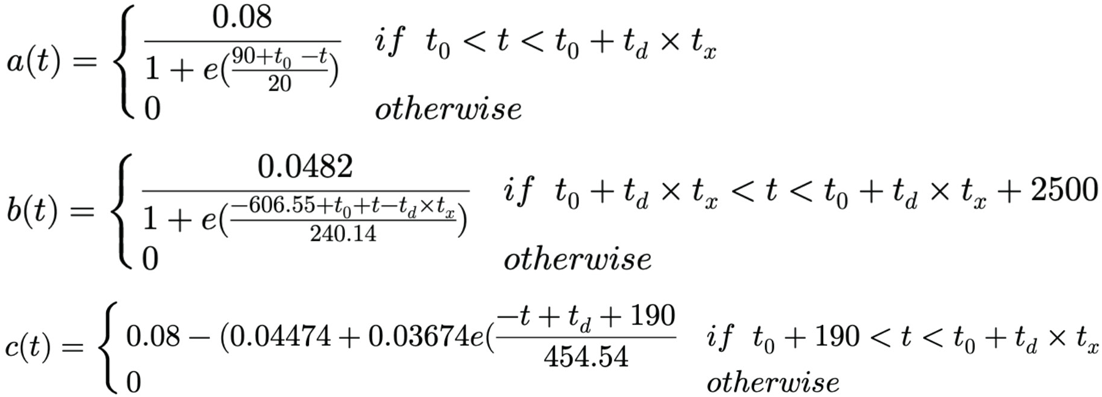

Where *t_0_* is the odour stimulus onset in ms, *t_d_* is the stimulus duration in ms and *t_x_*is a duration multiplier to reflect that receptor current outlasts the stimulus with this duration increasing with both the intensity and duration of the stimulus (*51–53*, *56*). For the weak stimulus *t_x_* was set at 1 and for the strong stimulus *t_x_*was 1.65 + a value drawn at random from a gaussian distribution with a mean of 0.2 and SD of 0.25 to reflect heterogeneity in the response decay across neurons carrying the same receptor (*77*). Peri-stimulus time histograms were computed for 500 olfactory receptor neurons at each concentration with bin widths of 50 ms. To estimate the Ca^2+^ signal that GCaMP6f would report for each odour concentration the mean spike rate was convolved with a kernel representing the kinetics of GCaMP6f (*21*).

## Acknowledgments

This work was funded by the MRC (MR/V003747/1) and by a studentship funded by the University of Leeds. We would like to thank Emma Smith who provided technical support to the project and Lucy Hurst and Mimi Levinson who performed initial key point assessment for head-fixed video analysis.

## Notes

### Competing Interest Statement

The authors have declared no competing interest.

### Summary of Updates

Substantial new data has been added which changes the focus of the manuscript

